# Signaling mechanism of phytochromes in solution

**DOI:** 10.1101/2020.04.05.025882

**Authors:** Linnéa Isaksson, Emil Gustavsson, Cecilia Persson, Ulrika Brath, Lidija Vrhovac, Göran Karlsson, Vladislav Orekhov, Sebastian Westenhoff

**Affiliations:** Department of Chemistry and Molecular Biology, University of Gothenburg, Gothenburg; Swedish NMR center, University of Gothenburg, Gothenburg

## Abstract

Phytochrome proteins guide the red/far-red photoresponse of plants, fungi, and bacteria. The proteins change their structure in response to light, thereby altering their biochemical activity. Crystal structures suggest that the mechanism of signal transduction from the chromophore to the output domains involves refolding of the so-called PHY tongue. It is currently not clear how the two other notable structural features of the phytochrome superfamily, the helical spine and a figure-of-eight knot, are involved in photoconversion. Here, we present solution NMR data of the complete photosensory core module from *D. radiodurans* (*Dr*BphP). Photoswitching between the resting and active states induces changes in amide chemical shifts, residual dipolar couplings, and relaxation dynamics. All observables indicate a photoinduced structural change in the knot region and lower part of the helical spine. This implies that a conformational signal is transduced from the chromophore to the helical spine through the PAS and GAF domains. The new pathway underpins functional studies of plant phytochromes and may explain photo-sensing by phytochromes under biological conditions.

## Introduction

Phytochromes are universal photoreceptors found in plants, fungi, and various microorganisms (Butler et al., 1959). They detect red and far-red light, thereby controlling many aspects of growth, development, and movement. The functions range from phototaxis and pigmentation in bacteria to seed germination, shade avoidance and flowering in higher plants (Yeh, 1997, Hughes et al., 1997, Davis, 1999, Fankhauser, 2001, Rockwell and Lagarias, 2017, Wagner et al., 2005, Franklin and Quail, 2010). Phytochromes work by switching between two photochemical states. In prototypical phytochromes, the resting state absorbs red light (Pr state), and the activated state absorbs far-red light (Pfr state). The proteins can be reversibly switched between the states with red light (Pr*→*Pfr) and far-red light (Pfr*→*Pr). Phytochromes are important for life, since they ensure that microbes and vegetation can adapt to light and thrive on earth.

Similar to other signaling proteins (Möglich et al., 2010, Möglich, 2019), phytochromes are modular proteins. They are divided into three groups according to the domain architecture. Group I phytochromes share a highly conserved photosensory module consisting of the domains PAS/GAF/PHY (Per/Arnt/Sim-cGMP phosphodiesterase/adenyl cyclase/Fh1Aphytochrome specific) (Auldridge and Forest, 2011, Rockwell et al., 2006). Plant, fungal and bacterial phytochromes belong to this group. The photosensory module of the cyanobacterial group II and III phytochromes consist of GAF/PHY and GAF, respectively (Anders and Essen, 2015). A covalently linked bilin chromophore (biliverdin in bacterial phytochromes) is attached via a thioether linkage to a conserved cysteine in the GAF domain (cyanobacteria and plants) or preceding the PAS domain (bacteria) (Auldridge and Forest, 2011). In bacteria, the output domains are often C-terminal histidine kinases. In plants or fungi, output domains consist of N-terminal extensions (NTE) and C-terminal PAS domains connected to a kinase-like domain (Rockwell et al., 2006). Phytochromes are usually homodimers.

The photosensory module of phytochromes contains three characteristic structural elements (numbering throughout the paper is for *Dr*BphP). These are a conserved loop region in the PHY domain, called “tongue” (residue 444-476, group I and II), which extends from the PHY domain back towards the chromophore binding pocket, a figure-of-eight “knot” in the PAS/GAF domain (residue 27-38 and 228-256, group I), and a long helix called “spine” (residue 300-345, groups I-III), which extends through the entire photosensory module (Fig. 1A). The spine helix is coupled to another helix in the PHY domain, which forms a coiled-coil with the equivalent helix in the sister-monomer connecting the photosensory module to the output domain (Gourinchas et al., 2017, Otero et al., 2016).

**Figure 1:**
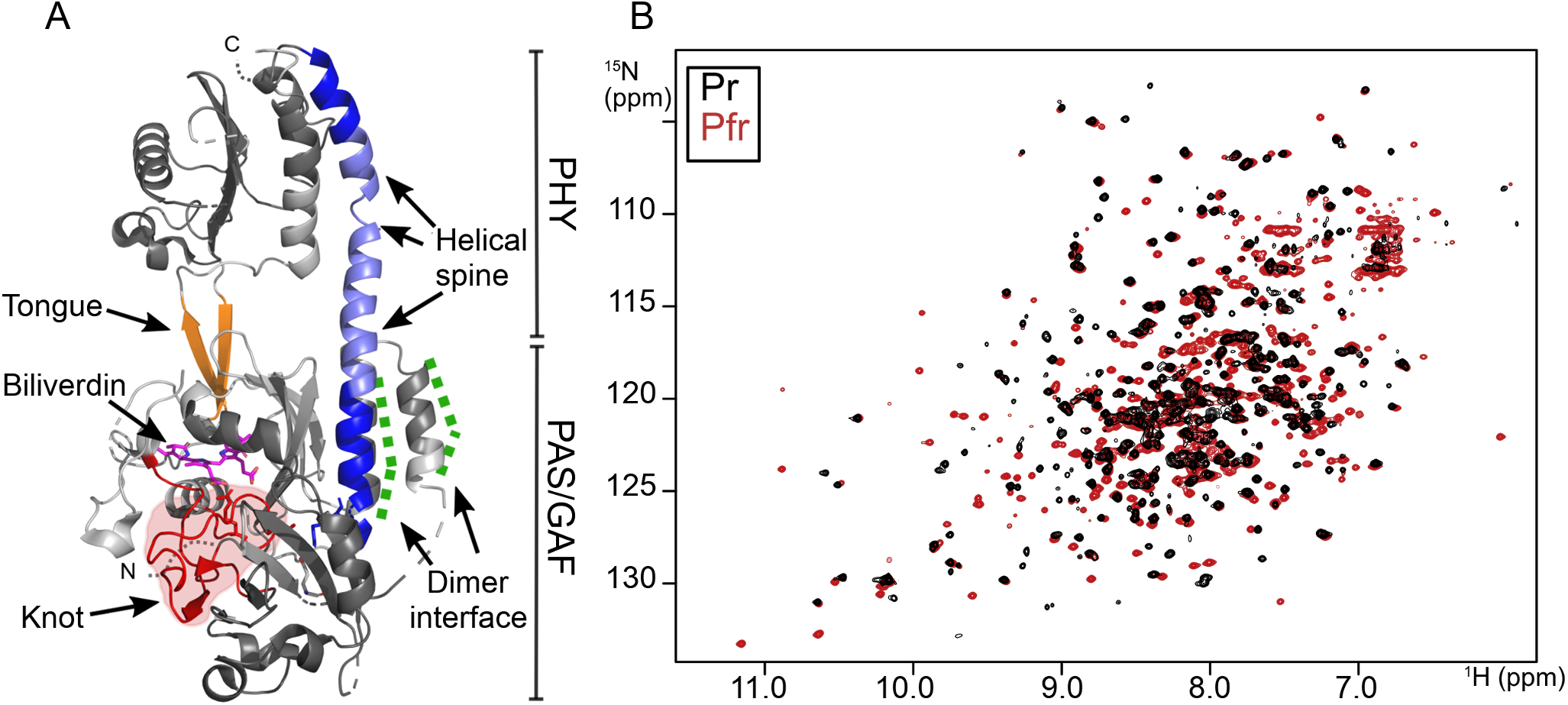
NMR characterization of *Dr* BphP . **A** The structure of the Pr state photosensory module of *Dr* BphP (PDB ID: 4O0P) is displayed with the tongue (orange), knot (red), biliverdin (magenta), and helical spine (blue) indicated. The lighter colors (marked light blue/light grey) mark residues which have not been assigned in both states. The middle part of the **B**2D [^1^H^15^N]-TROSY spectra for the dark (Pr) and light (Pr/Pfr mixture) state are displayed. The assignment for the Pr and Pfr states was obtained in Ref. (Gustavsson et al., 2020).

Crystal and solution structures of phytochromes in Pr and Pfr states indicate a movement of the PHY domains with respect to the PAS/GAF domains (Takala et al., 2014, Burgie et al., 2016, Assafa et al., 2018, Takala et al., 2016). How this change arises is currently debated. The tongue refolds from a *β*-hairpin conformation in Pr to an *α*-helical structure in Pfr (Takala et al., 2014). Interestingly, crystal structures have indicated a much quieter response to photoactivation in the knot and in the helical spine region. As a result, the tongue is currently considered to be the main signaling route in phytochromes (Takala et al., 2014, Anders et al., 2013). This is not fully satisfying given that many fully functional phytochromes exist, which lack the tongue (Anders and Essen, 2015). Therefore, another signaling pathway may exist.

The knot in group I phytochromes is formed at the interface between the PAS and GAF domains spanning the space between the chromophore and the helical spine (residue 27-38 and 228-256) (Fig. 1A) (Wagner et al., 2005). The sequence between Cys24 and the start of the PAS domain (residue 27-38) is passed through a loop formed by a large insertion within the GAF domain (residue 228-256) (Wagner et al., 2005). The knot is stabilized by a small but critical hydrophobic core, centered on the conserved Ile35. The region contains many conserved residues (Ile35, Gln36, Leu41, Ala225, Val232, Leu234, Leu248, Leu253 and Arg254) corroborating its importance for the protein (Wagner et al., 2005). It has been speculated that the knot stabilizes contacts between the PAS and GAF domains, or that it restricts movement of the N-terminal domain and thereby orients Cys24 for efficient conjugation to biliverdin. Another theory is that the knot limits the flexibility of the protein in order to prevent undesirable loss of entropy due to large domain movements upon photoisomerization of the chromophore (Wagner et al., 2005). However, as current crystal structures do not reveal any changes in the knot region between Pr and Pfr (Otero et al., 2016, Burgie et al., 2016, Takala et al., 2014, Essen et al., 2008, Yang et al., 2008, Takala et al., 2018, Schmidt et al., 2018), these ideas remain speculative.

The helical spine consists of two long helices, one connecting the PAS/GAF domain with the PHY domain (’lower’ spine helix, Fig. 1A) and one connecting the PHY to the output domains (’upper’ spine helix). Helical spines are common in many signalling proteins and they are actively involved in signaling (Möglich, 2019, Gushchin et al., 2017, Bhate et al., 2015). This also seems likely for phytochromes and has been suggested when the first crystal structures of photosensory domains were disclosed (Yang et al., 2008, 2009). Indeed, photoactivation causes a bend of the spine helix between the PAS/GAF and PHY domains as inferred from crystal structures, X-ray solution scattering, and electron paramagnetic resonance spectroscopy (*SI Appendix,* Fig. S2A) (Takala et al., 2014, Burgie et al., 2016, Assafa et al., 2018, Takala et al., 2016, Bjorling et al., 2016). This is reasonable, as the spine provides a scaffold for the protein. However, it remains unclear if the spine helix has a more active role in transducing the photosignal towards the output domains.

In order to uncover the signal transduction mechanism, the protein should be studied in solution. Solution Nuclear Magnetic Resonance (NMR) is an ideal tool to characterize dynamic molecules at atomic resolution. Chemical shifts are very sensitive probes for locating and detecting conformational changes of amino acids or their environment (Williamson, 2013, Oschkinat et al., 1988). Residual dipolar couplings (RDCs) provide information on the orientation of chemical bonds in relation to a molecular frame, defined by the protein. They provide long-range orientational information on the single-residue level (Bax and Grishaev, 2005, Lipsitz and Tjandra, 2004). T1 and T2 relaxation measurements give information on changes in structural dynamics between states. Of special interest is the T1/T2 ratio, which in a significantly non-spherical molecule indicates the relative orientation of each amide bond with respect to the molecular frame.

The investigation of phytochromes by NMR spectroscopy is difficult, because of their large size. Therefore, solution-state NMR has only been applied to short phytochrome constructs, which consist of single GAF domains (Lim et al., 2018, Cornilescu et al., 2014, 2008, Ulijasz et al., 2010), and residues around the labelled chromophore in PAS/GAF constructs (Strauss et al., 2005). However, the knotted PAS/GAF domain or the complete photosensory module have not been investigated, because they were considered to be too large (Lim et al., 2018). Solid state NMR has been used to study the chromophore binding pocket of larger phytochrome fragments (Song et al., 2011, 2018, 2014).

We have recently presented the first solution NMR assignment of the complete photosensory module of a uniformly triple-labelled (^2^H,^13^C,^15^N) phytochrome (Figure 1B, BMRB Entries 27783 and 27784 for Pr and Pfr states, respectively) (Gustavsson et al., 2020), using the monomeric version of the *D. radiodurans* bacteriophytochrome (PAS/GAF/PHY) (Auldridge et al., 2012, Takala et al., 2015) (*SI Appendix,* Fig. S1). 73% and 75% of assignable residues were assigned in Pfr and Pr, respectively. The assignments open up the opportunity for a detailed investigation of the photoinduced changes in *Dr* BphP_*mon*_ in the present study. We present chemical shift perturbation analysis, RDCs, and ^15^N T1 and T2 relaxation data for Pr and Pfr state for *Dr* BphP_*mon*_ . The observables describe a clear difference between the Pr and the Pfr state in the knot region and the lower part of the helical spine, identifying a new signaling route in phytochromes.

## Results

### Chemical shifts in Pfr and Pr indicate remodelling of the PAS and GAF domains

Fig. 2A shows the changes of the amide chemical shift between Pr and Pfr for *Dr* BphP_*mon*_, which we computed from the 2D [^1^H^15^N]-TROSY spectra (Fig. 1B). The analysis was performed for 231 residues that are assigned in both the Pr and Pfr state. Most residues show only small changes, but 47 residues show differences above 0.1 Δ*ppm* and 15 above 0.2 Δ*ppm*. Fig. 2B shows the location of these amino acids in the structure. Except for four residues, which are located close to the tongue region in the PHY domain, all other residues are found in the PAS/GAF domains, with a particular focus in the knot and helical spine regions (see zoom-in in Fig. 2B). In particular, residues ranges 33-40, 225-232, 251-257, and 300-311, have the largest changes in chemical shift and are located in the knot region and the lower part of the helical spine (Fig. 2A and B). Residue Phe61, which also has a notable change in chemical shift, is located close to the bottom part of the helical spine (Fig. 2B). The chemical shift perturbation analysis does not cover the tongue region and the middle part of the helical spine, because assignment is lacking in Pr or Pfr states (*SI Appendix,* Fig. S1) (Gustavsson et al., 2020). Chemical shift changes indicate chemical or structural changes in the near surrounding of the relevant amide groups. We therefore suggest that, in solution, previously unrecognized structural or chemical changes occur in the PAS/GAF domain.

**Figure 2:**
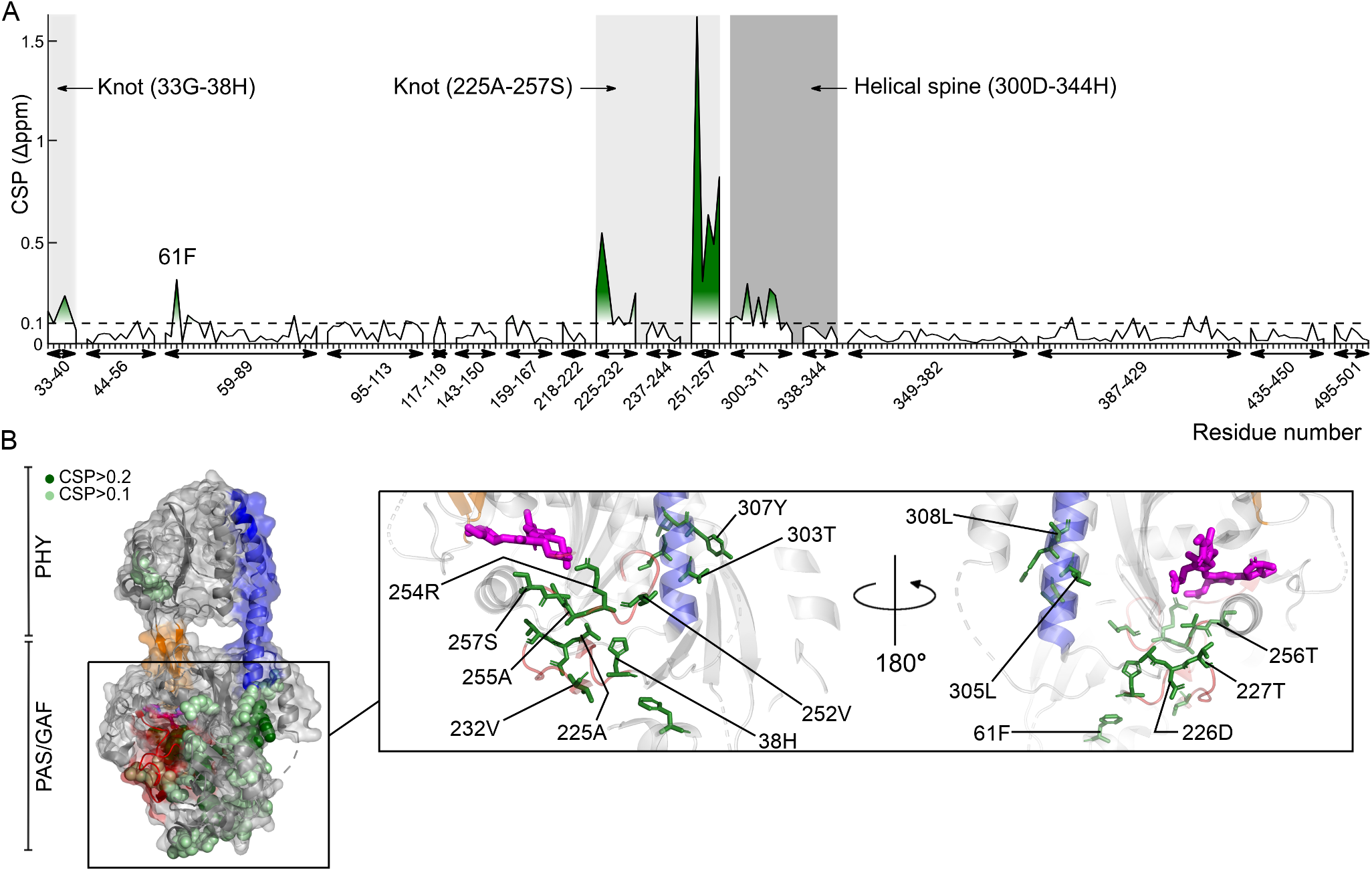
Chemical shift perturbation data for N and NH show large changes in chemical shifts in the knot and the helical spine. **A** The chemical shift perturbation (CSP) of the amide chemical shift is shown. The knot and lower helical spine are indicated. Only residues which are assigned in Pr and Pfr are displayed. For full set of CSP data, see *SI Appendix,* Fig. S6. The chemical shift perturbation was calculated as 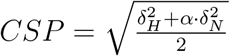 with *α* = 0.15. **B** The same structure as Fig. 1A of the photosensory module is shown with chemical shift changes indicated in green colors. Residues with CSP*>*0.2 are marked in dark green in zoom-in on the PAS/GAF domains. Leu253 unfortunately lacks assignment.

### The secondary structure remains unchanged in the knot and the spine region

Next, we calculated the secondary chemical shifts for CA (*SI Appendix,* Fig. S4A), CO (*SI Appendix,* Fig. S4B) and CB ((*SI Appendix,* Fig. S4C). These shifts collectively report on the secondary structure of the residues. We observed that the sign and the intensity for the secondary chemical shifts did not change between Pr and Pfr and we conclude that the secondary structure of the PAS, GAF, and PHY domains is retained in both states. We note that we cannot probe the secondary structure changes in the tongue region, because the tongue is not assigned in the Pr state due to structural heterogeneity in the Pr state(Gustavsson et al., 2020).

### Residual dipolar couplings (RDC) indicate changes in the knot and helical spine

RDCs are NMR observables, which report on structure. We measured RDCs on the amide N-H bonds using Pf1 phages for partial molecular alignment of the proteins.

To test if the Pr and Pfr crystal structures represent the solution structures, we inspected correlation plots between experimental RDCs for Pr and Pfr and RDCs computed from the crystal structures (4Q0J and 5C5K respectively) of *Dr* BphP_*PSM*_ using the software PALES (Fig. 3 and Table 1) (Zweckstetter and Bax, 2000, Zweckstetter, 2008). We performed the assessment for RDCs in the complete photosensory core (PAS/GAF/PHY), in the PAS/GAF domain, and for the knot and the spine region. We classified the agreement into three categories according to the Q-factor (Cornilescu et al., 1998): *Q <* 0.4 for excellent agreement (Chen and Tjandra, 2011), 0.4 *≤ Q <* 0.8 for intermediate agreement, and *Q ≥* 0.8 for poor agreement. All results are visually confirmed in Fig. 3, where the gradient reports on correlation.

**Table 1:** Summary of RDC analysis. **Structure:** The crystal structures used in the RDC analysis. **Number:** Number of experimental RDCs. **RDC:** The RDC set used in the analysis. **PCC:** The Pearson correlation coefficient indicates the linear correlation between observed and calculated RDCs without rescaling. **Q:** The Q factor 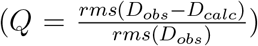 is an estimate of the average disagreement between observed and calculated RDCs (Cornilescu et al., 1998, Chen and Tjandra, 2011). See main text for classification of the “agreement” column.

| Structure | Number | RDC | PCC | Q | Agreement |
| --- | --- | --- | --- | --- | --- |
| Pr (4Q0J) | 176 | PAS/GAF/PHY (Pr) | 0.85 | 0.653 | intermediate |
| Pr (4Q0J) | 94 | PAS/GAF (Pr) | 0.93 | 0.361 | excellent |
| Pr (4Q0J) | 21 | Knot/Spine (Pr) | 0.945 | 0.248 | excellent |
| Pfr (5C5K) | 190 | PAS/GAF/PHY (Pfr) | 0.762 | 3.093 | poor |
| Pfr (5C5K) | 105 | PAS/GAF (Pfr) | 0.764 | 4.484 | poor |
| Pfr (5C5K) | 36 | Knot/Spine (Pfr) | 0.757 | 4.747 | poor |

**Figure 3:**
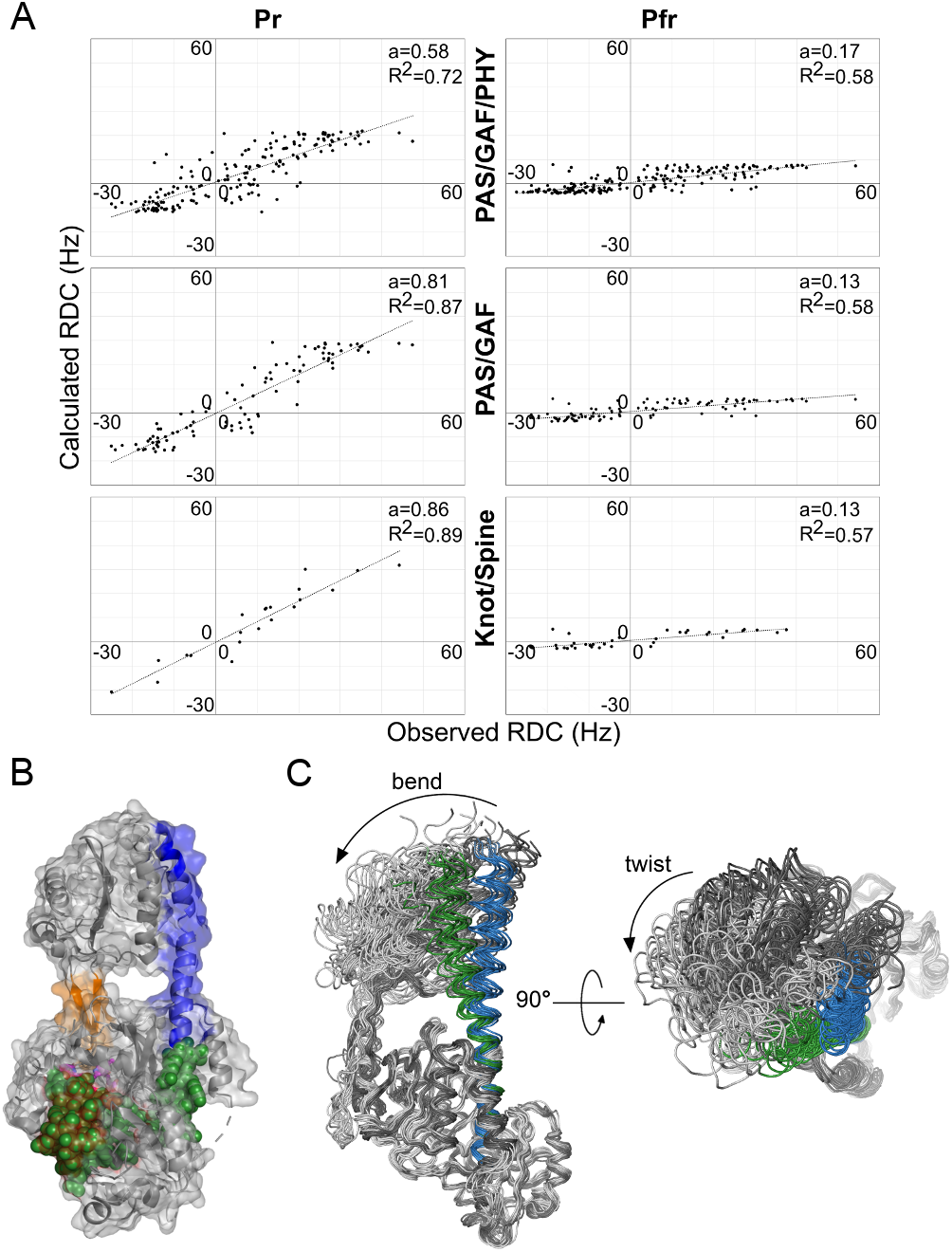
RDC data indicate changes in the PAS/GAF domains of *Dr* BphP . **A** Calculated RDCs are plotted against observed RDCs for Pr and Pfr. We used the PDB code 4Q0J for the Pr state calculations and 5C5K for Pfr state calculations. A linear fit to the data is shown with *a* as the gradient and *R*^2^ for the goodness of the fit. The computations were run for different fragments of the protein as indicated. **B** The color code in the structure is as in Fig.2 and the knot and spine region (residues 27 to 38, 228 to 256 and 299 to 344) are marked in green. **C** The overlay of the Pr and Pfr structures refined against the RDC data. The 10 best fitting structures for Pr (dark grey/blue) and Pfr state (light grey/green) are shown. All structures are aligned on the PAS/GAF domains.

For Pr, we find intermediate agreement for PAS/GAF/PHY and excellent agreement for PAS/GAF and the knot/spine region. This indicates that the crystal structure for the PAS/GAF domain resembles the solution structure well, but that adding the PHY domain causes some discrepancy. We attribute this to floppiness of the PHY domain with respect to the PAS/GAF domain. This is expected from solution X-ray scattering (Björling et al., 2016), molecular dynamics simulations (Takala et al., 2015), and the variations in the position of the PHY domain in different crystal structures of the photosensory core module (Takala et al., 2014, Burgie et al., 2016, Essen et al., 2008, Yang et al., 2014).

For Pfr, all fragments yield poor agreement. This means that the solution structure of the PAS/GAF domain in Pfr diverts from the crystal structures (Wagner et al., 2005, Takala et al., 2014, Burgie et al., 2016), implying that the crystal structures have not captured the structural change that occurs in the PAS/GAF domain. Using the RDCs, we track the origin of this disagreement down to the knot/spine region, where poor correlation is observed in Pfr (Fig. 3 and Table 1). We conclude that, upon photoconversion in solution, structural changes occur in the helical spine and the knot region leading to a rearrangement of the PAS/GAF domains.

Protein structure changes can be refined against RDC data. To do so, we used a large number of candidate structures of the photosensory core module in Pr and Pfr generated by molecular dynamics simulations (Takala et al., 2016). We then computed the RDC values for each candidate structure using PALES (Zweckstetter, 2008). By comparing the correlation between computed and measured RDC values, we found that many candidate structures show good agreement with the RDC data for Pr (*SI Appendix,* Fig. S5). For the Pfr data, most candidate structures do not show any correlation, as expected. However, a few structures showed improved agreement. When the 10 best-fitting structures for Pr and Pfr are drawn together, a small bend and twist of the PHY domain with respect to the PAS/GAF domain becomes clearly visible (Fig. 3C). The change is in excellent agreement to what was refined from solution X-ray scattering data for *Dr* BphP_*P* *SM*_ and similar to the changes of the PHY domain refined by electron paramagnetic resonance spectroscopy for Cph2 from Synechocystis sp. PCC 6803 (Assafa et al., 2018, Takala et al., 2016). The presented analysis is limited by the amount of candidate structures and because RDC data from only one alignment medium was used. Also, the ‘best’ fits for Pfr are still quite poor, indicating that not all structural changes are covered by the best models. We believe that these undetected changes are in the PAS/GAF domain. Despite these limitations, the initial analysis of the RDC data confirms the displacement of the PHY domain in the photosensory core fragments of phytochromes (Takala et al., 2014, Burgie et al., 2016, Assafa et al., 2018, Takala et al., 2016). The analysis method is a promising avenue to refine more detailed structural changes in the knot and helical spine region, when more RDC data becomes available.

### Relaxation dynamics indicate structural changes in the knot/spine region

The rotational correlation time of the entire protein (*τ_c_*) is related to its molecular size and shape. *τ_c_* was calculated from the average ratio of ^15^N T1 (longitudinal) and ^15^N T2 (transverse) relaxation times. They were similar for the Pr and the Pfr states: 21.9*±*3 ns and 22.2*±*3 ns, respectively. This indicates that protein does not dimerize in a light-dependent manner and that the quaternary structure of *Dr* BphP_*mon*_ is not changed dramatically upon photoactivation, which is consistent with the time-resolved solution scattering of the same protein fragment (Takala et al., 2016).

We also determined ^15^N T1 and T2 relaxation times for each residue (Fig. 4B and C and *SI Appendix,* Fig. S3). Similar to RDC measurements, the photoinduced change of the T1/T2 ratio for each residue indicates a structural reorientation of the amide bond in the molecular frame (Fig. 4D). Some residues, which have large differences in T1/T2 ratios (Fig. 4E) are located in the PHY domain and many cluster in the knot and spine region of the protein (Fig. 4A). One of the regions with large changes Ala255-Thr256-Ser257) corresponds well with the Val252-Ser257 amino acid stretch, where the largest changes in chemical shifts were observed (see Fig. 2A). For some other residues, significant changes are recorded in T1/T2 ratios, but not in chemical shifts. This is not surprising, since both observables respond differently to changes in the protein structure and dynamics. An example is the stretch from 338 to 344 in the PHY domain, which is located at the upper end of the helical spine. Intriguingly, and in agreement with the chemical shift data, the relaxation data indicate changes of many residues, which are located between the chromophore and spine helix (Fig. 4A).

**Figure 4:**
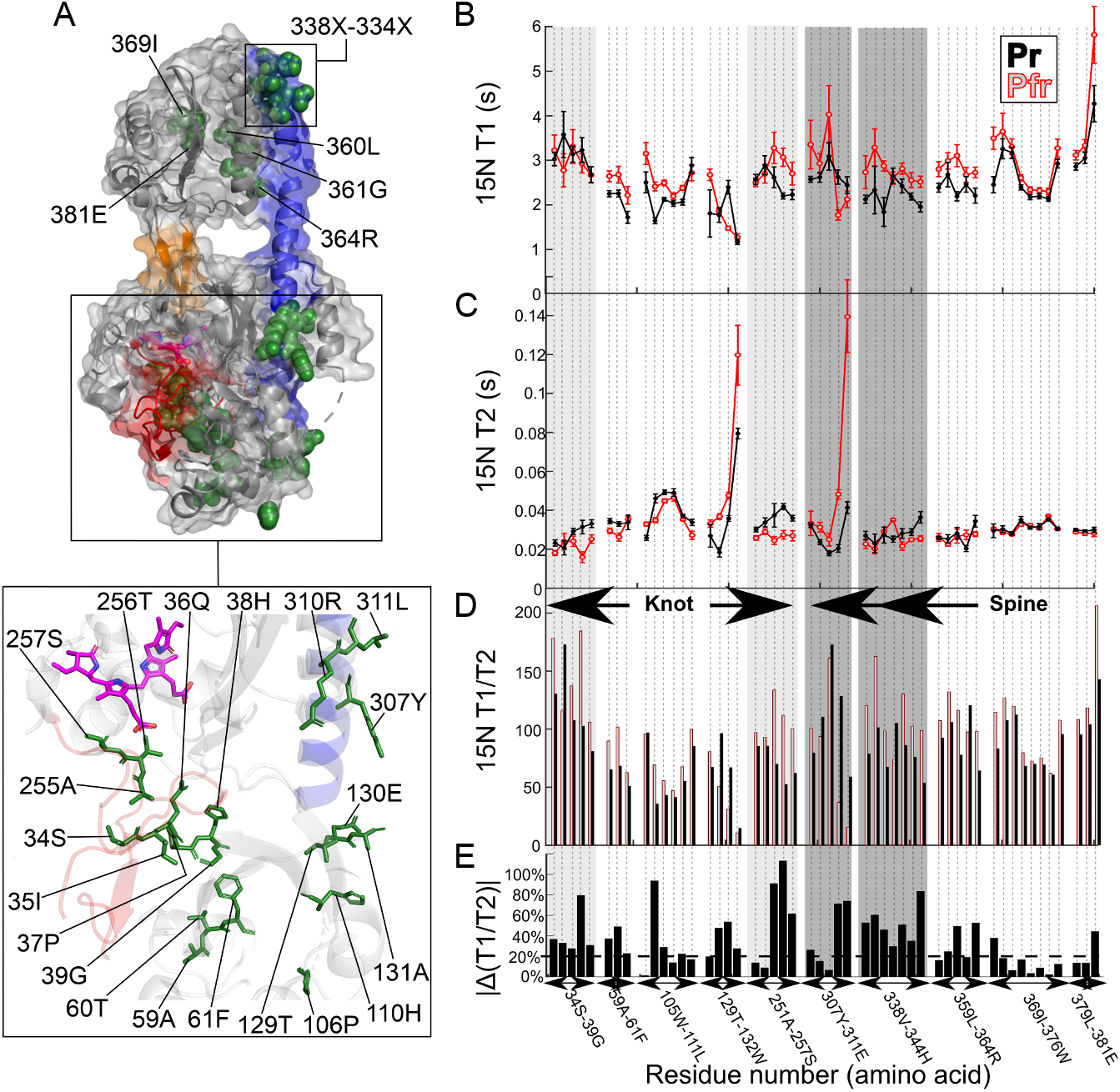
Large differences in the ^15^N T1/T2 ratios of the relaxation times indicate structural changes in the knot and spine region. **A** shows the *Dr*BphP structure (pdb ID: 4O0P) with the residues with large changes in the ^15^N T1/T2 ratio marked in green. The other colors mark structural elements as in Figure 1. **B** displays the ^15^N T1 relaxation, **C** the ^15^N T2 relaxation and **D** the ^15^N T1/T2 ratio for Pr (black) and Pfr (red). In **E**, the change of the T1/T2 ratio is shown in percent of T1/T2 of the Pr state. Changes larger than 20% (dashed line) where mapped on the structure in **A**.

### The NMR observables collectively reveal a signaling route through the knot and spine region

The analysis of chemical shifts, RDCs, and T1/T2 ratios indicate that significant structural changes occur when *Dr*BphP is photoconverted from Pr to Pfr in solution. The changes in chemical shift are residue-specific and they pinpoint changes to the knot region and the helical spine (Fig. 2C). RDCs report directly on the orientation of the amide bonds. They indicate structural changes of the PHY domain with respect to the PAS/GAF domains and a *structural* perturbation of the protein in knot and spine region, similar to the chemical shifts (Fig. 3). Moreover, the RDCs show that the crystal structures in Pfr do not represent the solution structure well, in particular in the PAS/GAF domain. T1/T2 is also indicative of structural changes, further confirming these changes (Fig. 4). The secondary structure is not changed, precluding unfolding of the domains. Nevertheless, we conclude that significant photoinduced structural changes arise in the PAS/GAF domains and that these changes occur in the knot region and the helical spine.

Fig. 5 illustrates this new proposed signaling pathway. The NMR data indicate alterations spanning from the chromophore to the helical spine. Arg254 (NMR data: CSP=0.31) is connected to the chromophore and is the entry point to the signal transduction chain. In the Pr state, Arg254 has a salt bridge to the propionate group of the B ring of the biliverdin (Takala et al., 2014, Essen et al., 2008, Burgie et al., 2014), which is broken in Pfr (Burgie et al., 2016, Song et al., 2011). The signal may continue through Val252 (NMR data: CSP=1.61), which is located next to the conserved Leu253 (lacking assignment), and via the conserved Glu127 (lacking assignment) that forms a salt bridge in the crystal structure to Arg302 (NMR data: CSP=0.12 and large T1/T2 difference). Arg302 is located in the bottom part of the helical spine. The notable change in chemical shift of Phe61 (NMR data: CSP=0.32) is consistent with this proposed structural change, as the residue is located within 10Å of the salt bridge. Many other neighbouring residues show changes in chemical shift and T1/T2, supporting the notion that the changes are more extensive. Taken together, the detected changes wire the chromophore to the lower part of the helical spine, constituting a potential signaling pathway. Given the similarity of phytochrome structures of the PSM in the PAS/GAF region (Otero et al., 2016, Burgie et al., 2016, Takala et al., 2014, Essen et al., 2008, Yang et al., 2008, Schmidt et al., 2018, Burgie et al., 2014), we propose that the signaling chain is widely conserved and that it is an integral part of the photoresponse of phytochromes.

**Figure 5:**
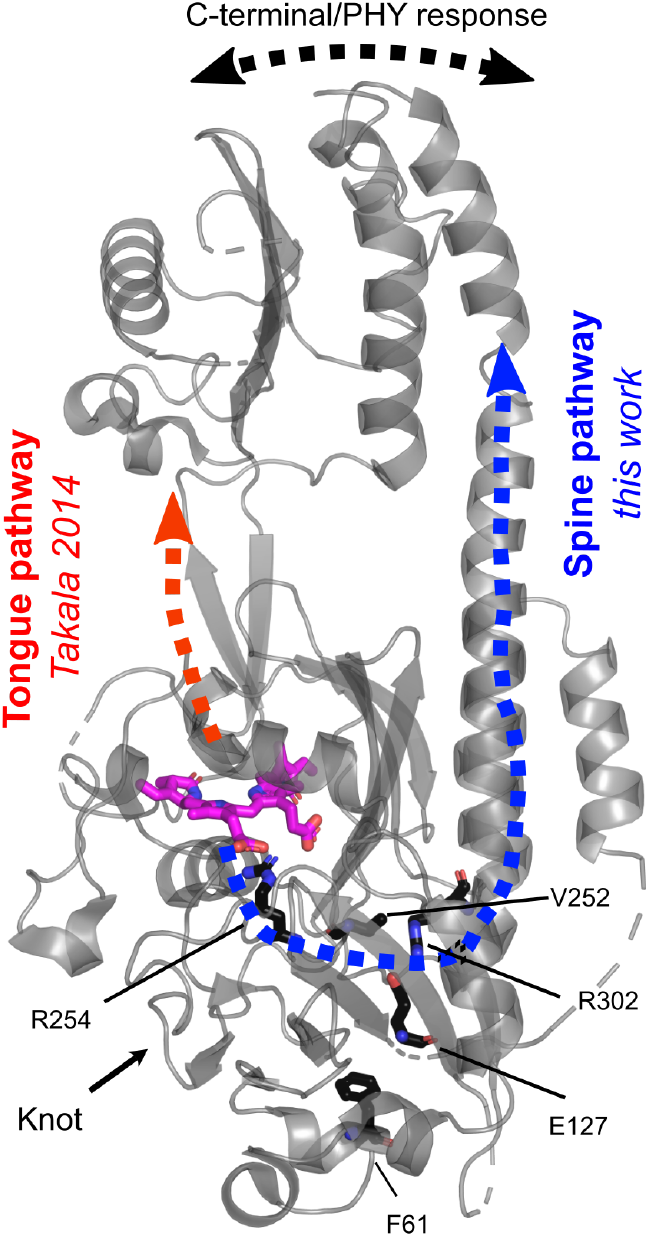
The proposed “spine” signaling pathway for phytochromes. The structure of *Dr*BphP (PDB ID: 4O0P) is shown together with the two pathways, which guide the signal to the PHY-domain and further to the C-terminal output domains. The “tongue” pathway (red) was proposed in 2014 based on crystal structures by Takala *et al.* and involved refolding of the tongue region (Takala et al., 2014). The new signaling pathway, which is proposed here with solution NMR spectroscopy, proceeds through the knot and spine region. Conserved amino acids, which participate in the pathway, are indicated.

## Discussion

It seems reasonable that all three outstanding and conserved structural elements of the phytochrome photosensory module are involved in the function of the protein. Here, we show that a signal transduction pathway exists, which spans from the chromophore via the knot to the lower part of the helical spine. Conceivably, these changes may translate to the C- and N-terminal output domains. Previous crystallographic work has only shown that the PHY-tongue is involved in the transduction of the light signal (Fig. 5), but changes in the knot and spine region inside the PAS/GAF domains were not observed (illustrated in *SI Appendix,* Fig. S2) (Wagner et al., 2005, Takala et al., 2014, Burgie et al., 2016, Assafa et al., 2018, Essen et al., 2008, Yang et al., 2008).

Generally, knotted polypeptide chains are expected to have an evolutionary rationale (Wagner et al., 2005, Dabrowski-Tumanski and Sulkowska, 2017). The phytochrome knot contains many highly conserved residues (Wagner et al., 2005). It has been demonstrated that the knot region is important for signaling in a plant phytochrome. Mutations R110Q, G111D, G112D, and R352K in *Arabidopsis* PhyB (corresponding to Ile31, Pro32, Gly33 and Arg254 in *Dr*BphP) have normal photochemistry, but are defective in intracellular signal transfer (Oka et al., 2008). The signal transfer is blocked, because binding to the Phytochrome Interacting Factor (PIF, basic helix-loop-helix transcription factors) is disrupted by the mutations (Kikis et al., 2009). One end of the knot is the start of the N-terminal extension (Fig. 2A), which is known to be of functional relevance in plant phytochromes (Nito et al., 2013, Trupkin et al., 2007, Medzihradszky et al., 2013, Cherry et al., 1992).

Even the somewhat shorter N-terminal segment in bacterial phytochrome changes between crystal structure in Pr and Pfr (Essen et al., 2008, Yang et al., 2008, Gourinchas et al., 2019) and influences the spectral properties (Gourinchas et al., 2019). In light of these findings it is conceivable that the knot is actively involved in signal transduction.

Helical spines are common structural elements in many signaling proteins and critical for signal transduction (Möglich, 2019, Gushchin et al., 2017, Bhate et al., 2015, Berntsson et al., 2017). While the helical spine exists in all phytochromes, the PHY-tongue does not, because PHY-less phytochromes exist (Auldridge et al., 2012). Coiled-coil interactions between the C-terminal extension of the helical spine and the output domain have been shown to be critical for controlling the signaling output in phytochromes (Gourinchas et al., 2018), resembling the converged view for other bacterial histidine kinases (Möglich, 2019, Bhate et al., 2015). These considerations provide support for our proposed new pathway for signal transduction from the chromophore via the knot region to the lower part of the helical spine.

With the presented NMR experiments, we were not able to characterize structural changes in the tongue region, because the tongue is not assigned in the 3D TROSY-type H-N-C spectra in the Pr state. However, based on the crystallographic evidence (Takala et al., 2014, Burgie et al., 2016), we assume that this region changes upon photoactivation. Even the NTE region, which interacts with the tongue region, is likely to change,Gourinchas et al. (2019) but is unfortunately unassigned.(Gustavsson et al., 2020). As such, there would be two pathways for signal transduction in the (PAS/GAF/PHY) photosensory module of phytochromes; one through the PHY-tongue (Takala et al., 2014) and another one through the knot and helical spine. It is difficult to say which part of the protein is changed first. We consider that the two signaling routes may work in concert to stabilize each other, or that they are redundant signaling pathways.

An overlay of the Pr and Pfr structures for *Dr* BphP_*P* *SM*_ shows virtually no change in the PAS/GAF domain (Takala et al., 2014, Burgie et al., 2016, Essen et al., 2008, Yang et al., 2008), but now our NMR data indicate substantial changes in these domains. Our RDC data show, that the crystal structure resembles the solution structure well in Pr, but not in Pfr. How can this be rationalized? It is known that the photophysical state of the chromophore can be uncoupled from the conformation of the phytochrome in crystal structures (Otero et al., 2016, Takala et al., 2018). The energy of formation of crystal has the same order of magnitude as the energy of interactions between protein subunits. Because these energies are similar, crystal packing can alter protein conformations. It could be that the Pfr conformation of PAS/GAF in solution has a disordered connection between PAS and GAF domains. Such a ‘loose’ conformation would not be favoured in tightly packed crystals. It could also be that one of the conformations causes a disruption of the dimer packing, which would also not be preferred in crystals. The unnatural situation of tight packing in crystals then pushes the protein back into the “Pr-like” conformation. We note that a difference signal (Pfr minus Pr) was recorded with solution x-ray scattering for the PAS/GAF fragment of *Dr* BphP (Björling et al., 2016). This signal indicates structural changes in the PAS/GAF domains in solution in agreement with the present NMR data.

Multiple conformations have been observed in the chromophore binding regions of phytochromes. Solid-state and solution NMR on the sensory module of Cph1 phytochrome and a GAF fragment from a red/green cyanobacteriochrome has revealed two isoforms of the red-light absorbing Pr state (Lim et al., 2018, Song et al., 2011). Double conformations have never been observed in crystal structures of phytochromes, emphasizing the detrimental effect that crystal packing can have. Even though we do not detect multiple conformations in the backbone chemical shifts near the chromophore, the presence of such heterogeneity is not in conflict with our model of signal transduction.

Although the collective NMR observables pinpoint the structural changes to the knot and spine region of the PAS/GAF domain (Fig. 2, 3, 4) they do not reveal an atomistic model of structural changes. One plausible structural mechanism is that the spine helix rotates and with that transduces the signal to the PHY and output domain. This would be consistent with structural changes, refined against time-resolved X-ray scattering data for *Dr* BphP_*mon*_ and for the full length *Dr*BphP (Takala et al., 2016, Björling et al., 2016). The studies identified a twist and bending of the photosensory core (Takala et al., 2016) and an overall twist of the output domains with respect to PAS/GAF (Björling et al., 2016). A rotation of the spine helix could also rationalize the bends or kinks that have been observed in the spine helix in crystal structures and by electron paramagnetic resonance for several phytochromes (Gourinchas et al., 2017, Burgie et al., 2016, Assafa et al., 2018, Takala et al., 2014, Essen et al., 2008, Yang et al., 2008, Burgie et al., 2014), and it would be consistent with how other sensory proteins are thought to work (Gushchin et al., 2017, Berntsson et al., 2017).

An alternative mechanism could be that the changes in the lower part of the spine helix lead to a modification of the dimer interface. The PAS/GAF part of the helical spine forms a significant fraction of the dimer interface. A rotational or translational motion could change the interactions across the dimer, thereby changing the dimeric arrangement. This would lead to modification of the coiled-coil connector from the PHY domain to the output domain, as proposed (Gourinchas et al., 2018). Indeed, the PAS/GAF fragment of *Dr*BphP crystallizes with different dimer arrangements in different crystal forms (Wagner et al., 2005, 2007, Edlund et al., 2016) and phytochromes typically have different dimerization constants in Pr and Pfr (Takala et al., 2014, Strauss et al., 2005).

To conclude, we propose a previously unrecognized signaling pathway along the knot and the spine helix in *Dr*BphP (Fig. 5), connecting the chromophore and the output domains. This pathway most likely works in concert with other pathways, for example along the PHY tongue. The work opens up the prospect for a more detailed detailed investigation of phytochromes with NMR spectroscopy, which we hope will refine the signal transduction cascade in the protein superfamily with atomic precision.

## Online Methods

Detailed information for the cloning, protein expression, protein purification, NMR sample preparation, NMR experiments, and data analysis can be found in *SI Appendix*.

## Acknowledgement

We would like to thank Markus Zweckstetter for feedback of the manuscript and support regarding the PALES software and Irena Matečko Burmann and Björn Burmann for critically reading the manuscript. Sebastian Westenhoff thanks the Knut and Alice Wallenberg Foundation for an Academy Fellowship. Vladislav Orekhov thanks the support by the Swedish Research Council (Research Grant 201504614). NMR spectroscopy was carried out at the Swedish NMR Centre at University of Gothenburg, supported by the Knut and Alice Wallenberg Foundation (NMR for Life) and the Science for Life Laboratory (SciLifeLab).

## Author contributions

LI, EG and SW designed the project. LI and EG planned the experiments together with CP, UB, GK, SW, and VO. LI and EG produced and purified all samples and performed the measurements and analysis together with CP, UB, VO, and SW. LV performed the MD analysis of the RDC data. LI, EG, VO and SW wrote the manuscript with input from all authors.

