## Supplemental information for "Signaling mechanism of phytochromes in solution" (from another version of this preprint: version 1, 2020-04-05)

### SI Appendix

### Material and methods

#### Cloning

The construct for the *Deinococcus radiodurans* monomeric phytochrome PAS/GAF/PHY fragment (57 kDa) has previously been cloned and transformed into *E.coli* BL21(DE3) cells.[1]

#### Expression and Purification

##### Triple-labeled phytochrome

Cells were adapted to $D_2O$ by streaking out on LB-agar plates containing increasing concentration of $D_2O$ (25, 50, 75, 100%). Colonies from the 100% $D_2O$ plate were picked and 5 ml overnight cultures of the phytochrome were prepared in M9 media ($Na_2HPO_4$, $KH_2PO_4$, NaCl, $^{15}NH_4Cl$, u-$^{13}$C-d-glucose, thiamine, $MgSO_4$, $CaCl_2$, $D_2O$). 1 ml of one of these cultures was inoculated to 50 ml of M9 media. This culture was further used to inoculate 2x500 ml of M9 media. At OD 0.6, the temperature was decreased to 30°C and 1 mM IPTG was added. Induction continued for 22 hours before the cells were harvested. An Emulsiflex C3 (Avestin) was used to disrupt the cells followed by a 30 minute centrifugation at 10 000 rpm. 3 mg of biliverdin (BV) hydrochloride (Frontier Scientific) was dissolved in 1 ml 30 mM Tris buffer pH 8.0 with 8M NaOH titrated until BV was completely solubilized. The cell lysate and the BV was incubated dark overnight at 4°C. The sample was filtered and loaded onto a 5 ml HisTrap HP column (GE Healthcare). The protein was eluted with a linear gradient of imidazole with 30 mM Tris, 50 mM NaCl, 300 mM imidazole pH 8.0. The protein was concentrated with a 30 kDa MWCO Vivaspin (Sartorius) to 3 ml and loaded to a HiLoad 16/600 Superdex 200 pg column (GE Healthcare) equilibrated with 30 mM Tris pH 8.0. Eluted fractions were concentrated as described above. Buffer was exchanged by centrifugation to 25 mM NaPi, 50 mM NaCl, 1 mM EDTA pH 6.9. UV-Vis was used to verify that the protein could photoswitch, and the purity was checked using SimplyBlue Coomassie (Invitrogen) staining of Mini-PROTEAN TGX Stain-Free gel (Biorad). PageRuler Prestained

Protein Ladder (Fermentas) was used as a standard.

#### NMR Sample Preparation

##### Triple-labeled phytochrome

0.8 mM of [u-$^{2}$H,$^{13}$C,$^{15}$N]-labeled phytochrome sample was mixed with 10% $D_2O$, 1x Complete EDTA-free (Roche) and 0.02% azide. A 3 mm shigemi tube was used. To measure the phytochromes in the Pr state, the sample in the NMR tube was illuminated in darkness using a 780 nm LED for 10 minutes. The sample was injected into the magnet in darkness. To convert the sample to the Pfr state, a 660 nm LED-laser (400mW) was connected to an optical fiber with a diffuser tip (Laser components) which was inserted into the sample inside the 3mm shigemi tube. The sample was illuminated for 5 seconds every 20 minutes during data acquisition to continuously keep the sample in the Pfr state.

##### Residual Dipolar Couplings

[u-$^{2}$H,$^{13}$C,$^{15}$N]-labeled phytochrome was mixed to a final concentration of 0.6 mM with 10% $D_2O$, 1x Complete EDTA-free (Roche) and 0.02% azide. A 3 mm shigemi tube was used without the piston. Reference spectra, without alignment media, were recorded for the Pr and Pfr states. 10 mg/ml Pf1 phages (ASLA Biotech) was added and spectra were recorded for both states.

#### NMR Experiments

All NMR experiments were performed at 45°C on a Bruker spectrometer with Larmor frequencies of 800 MHz equipped with a triple resonance TCI cryoProbe. Backbone chemical shift assignment for both the Pr and the Pfr state have previously been reported,[2] where 75% of all assignable peaks were assigned for the Pr state and 73% for the Pfr state. Unassigned residues cluster in the core region and the helical spine of the protein, likely due to poor protein back-exchange.Using our illumination conditions (660 nm), the Pfr state is estimated to have an occupancy of 65%.

##### Secondary chemical shifts

The secondary chemical shifts $\Delta\,\delta = \delta$ - $\delta_{rc}$ were obtained by subtracting the chemical shift of each residue ($\delta$) with the random coil value for the same residue ($\delta_{rc}$). The random coil data was obtained from Wishart and Sykes.[3,4]

##### Residual Dipolar Couplings

Reference spectra, without alignment media, were recorded for the Pr and the Pfr state using ARTSY (TROSY) method.[5] After addition of Pf1 phages, the same experiment was repeated for the Pr and the Pfr state.

##### Relaxation

$^{15}$N T1 and T2 relaxation times were measured using previously described TROSY type experiments.[6,7]

#### Data analysis

The spectra were processed with *nmrpipe* software.[8] T1 and T2 relaxation data were analysed using *DASHA* program.[9] The RDCs were analysed with *PALES*, with the following parameters: Pales -bestFit -pf1 -mcStruct -map 1000. BestFit uses iterative least-squares minimization and singular value decomposition to find the alignment tensors, while mcStruct uses structural noise Monte-Carlo analysis over 1000 iterations to determine the uncertainty in alignment tensors.[10,11]

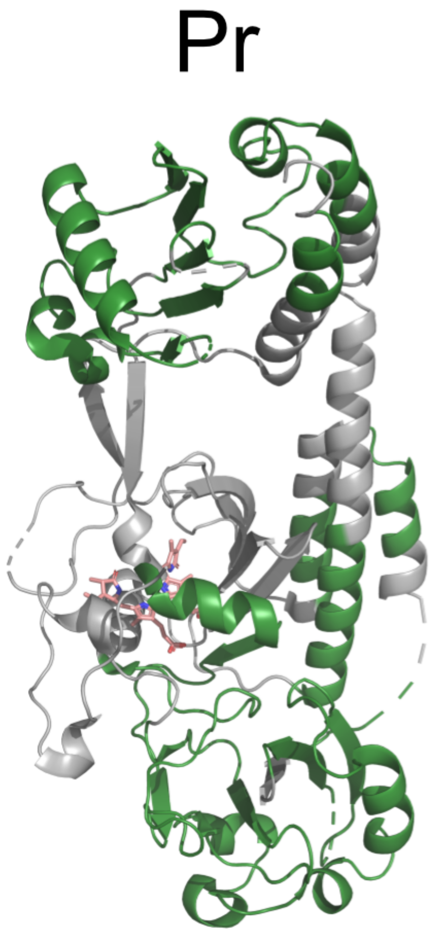


Figure 1: Backbone chemical shift assignment of the Pr (BMRB Entry 27783) and the Pfr (BMRB Entry 27784) state of the PAS-GAF-PHY photosensory module.[2] Assigned residues (green) are mapped on structures 4Q0J (Pr) and 4O01 (Pfr). The Biliverdin is coloured in pink.

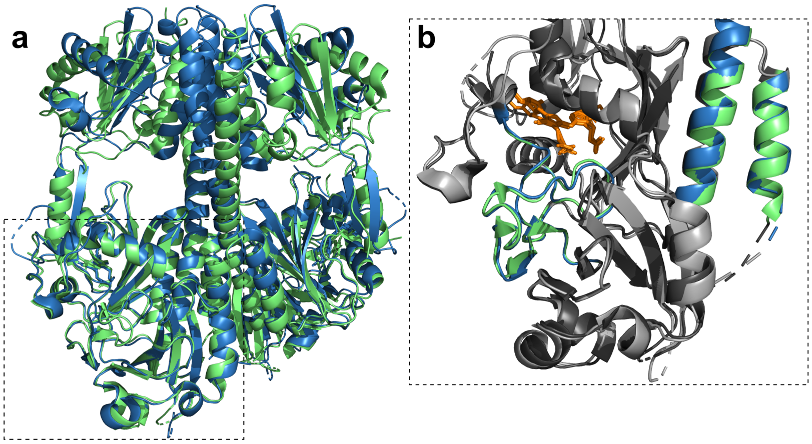


Figure 2: **a** Overlay of Pr (blue) and Pfr (green) structures, 4O0P and 4O01, respectively. **b** A zoom-in on the knot and dimer interface region (knot and dimer interface colored). No differences between Pr (blue) and Pfr (green) structures, 4O0P and 4O01, respectively, can be observed in the crystal structures for this region.

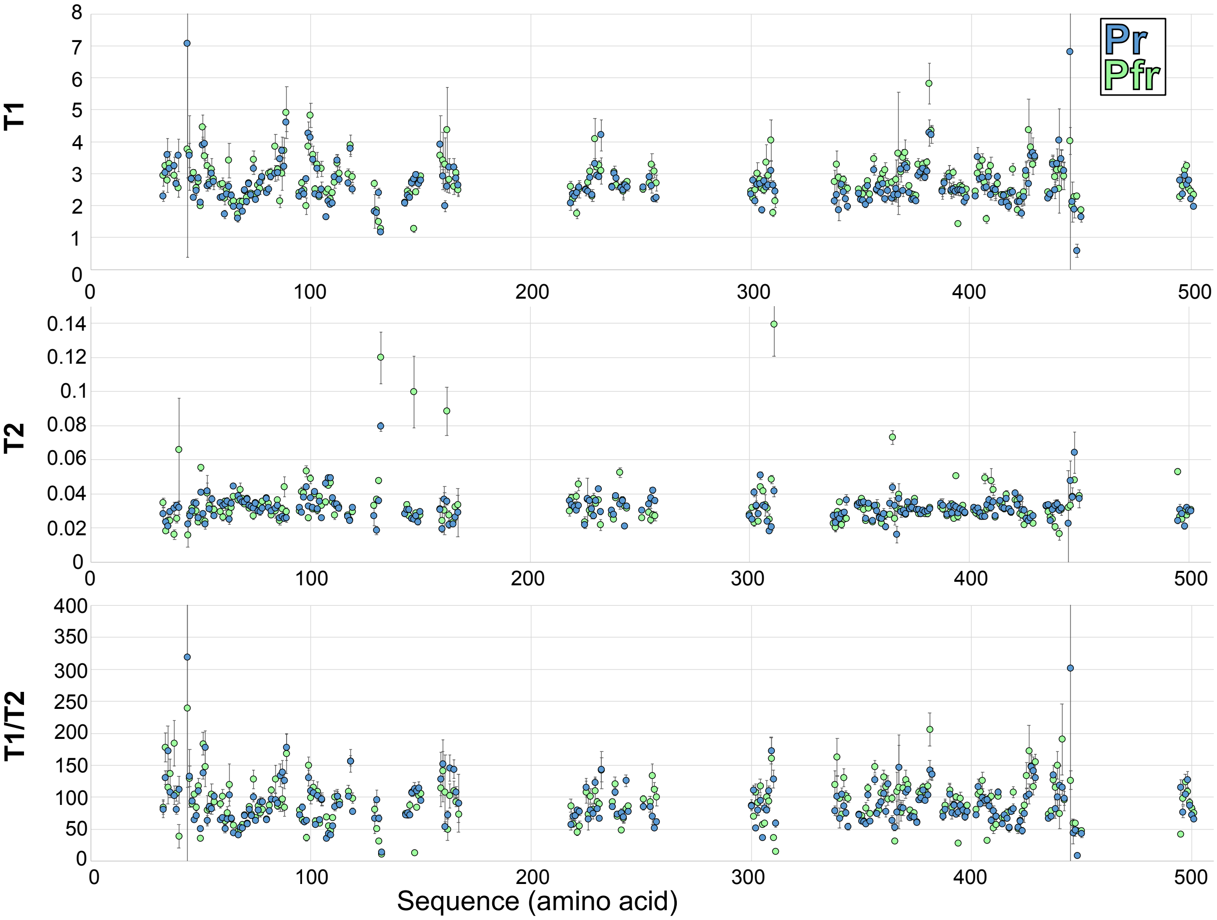


Figure 3: The complete data set of $^{15}$N T1, $^{15}$N T2 and $^{15}$N T1/T2 relaxation.

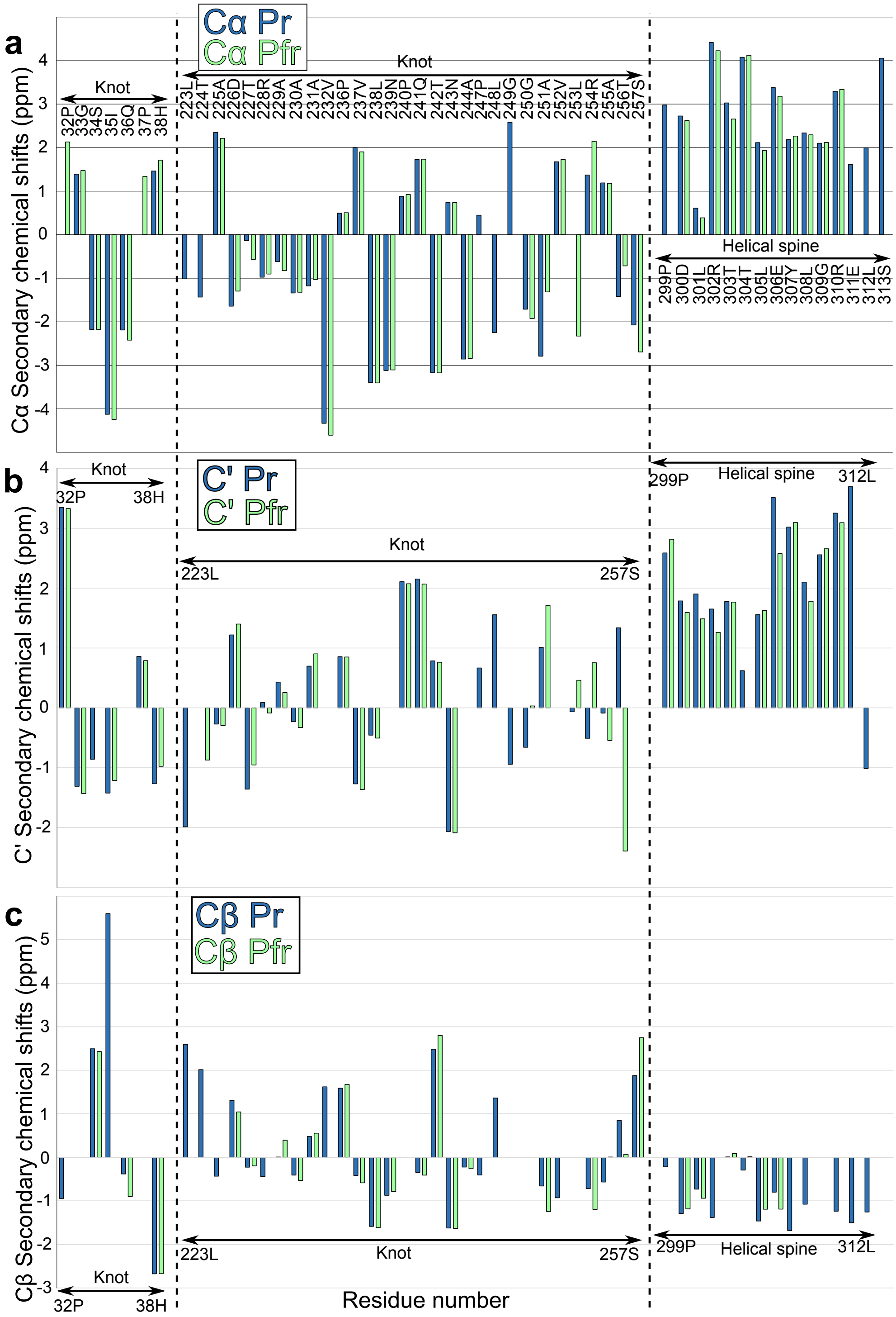


Figure 4: (**a**)C$\alpha$ secondary chemical shifts for the knot region and a part of the helical spine show no change between the Pr and the Pfr state. This excludes a secondary structure change in this region, which corresponds well with published crystal structures. (**b**) Secondary chemical shifts for C$\alpha$ (**a**), CO (**b**) and C$\beta$ (**c**). These secondary chemical shifts correlate well with the secondary structure elements in the crystal structures.

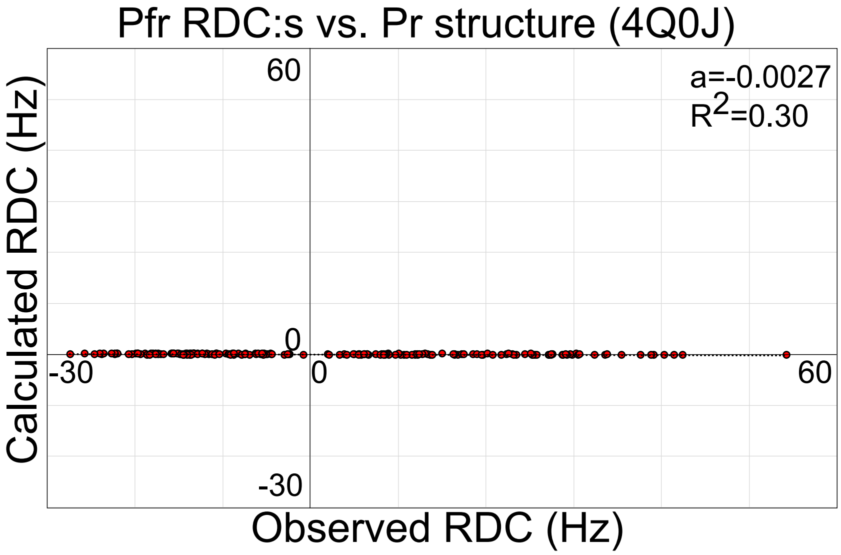


Figure 5: Experimental RDCs in Pfr compared to calculated RDCs from the Pr structure 4Q0J. Correlation is not observed.

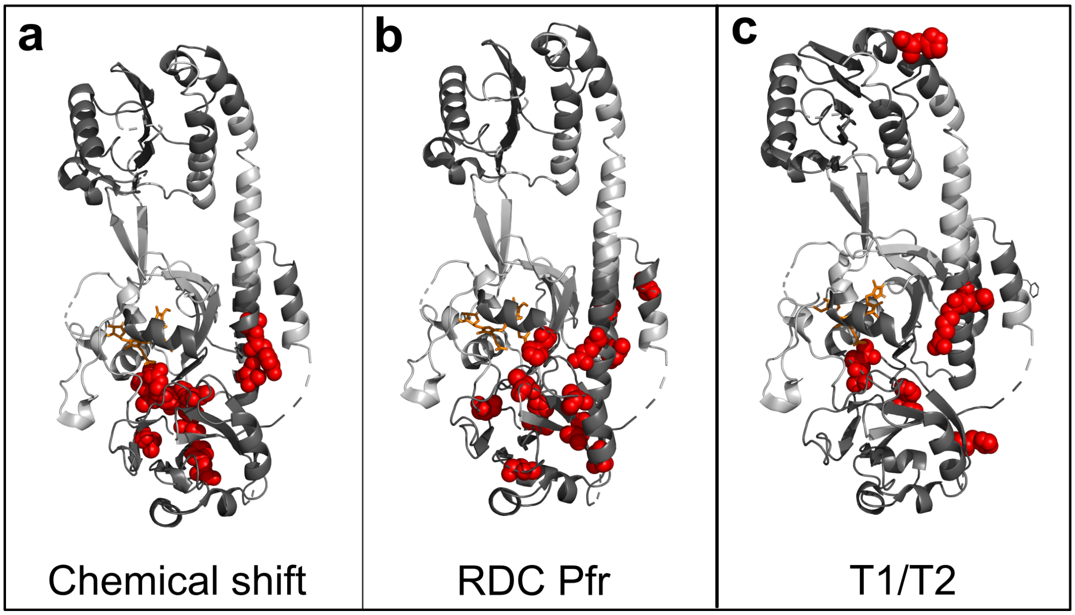


Figure 6: Overview of the observed changes from the three NMR experiments show that the knot and spine region are involved in signalling. PAS-GAF-PHY structure in the Pr state. Assigned residues are dark grey and unassigned light grey. THe biliverdin is marked in orange. **a** Large changes in chemical shifts (over $5\sigma$) are marked with red spheres. **b** The largest difference in $^{15}$N T1/T2 ratio between the Pr and Pfr states are marked with red spheres. **b** The largest RDC differences between the Pr and Pfr states are marked with red spheres.

Chemical Shift Perturbation

| Amino acid | CSP | Amino acid | CSP | Amino acid | CSP | Amino acid | CSP | Amino acid | CSP |
|---|---|---|---|---|---|---|---|---|---|
| **33 Gly** | 0.1601 | **88 Ala** | 0.0247 | **225 Ala** | 0.2672 | **354 Asp** | 0.0398 | **408 Leu** | 0.1254 |
| **35 Ile** | 0.0991 | **89 Leu** | 0.0869 | **226 Asp** | 0.5470 | **356 Ala** | 0.0185 | **409 Gly** | 0.0306 |
| **36 Gln** | 0.1690 | **95 Asp** | 0.0585 | **227 Thr** | 0.3286 | **357 Leu** | 0.0081 | **410 Gln** | 0.0878 |
| **38 His** | 0.2360 | **96 Ala** | 0.0869 | **228 Arg** | 0.0921 | **358 Asp** | 0.0113 | **411 Leu** | 0.0146 |
| **39 Gly** | 0.1509 | **97 Leu** | 0.0847 | **229 Ala** | 0.1325 | **359 Leu** | 0.0236 | **412 Trp** | 0.0236 |
| **40 Ala** | 0.0675 | **98 Gln** | 0.1064 | **230 Ala** | 0.0909 | **360 Leu** | 0.0612 | **414 Ala** | 0.0285 |
| **44 Ala** | 0.0228 | **99 Tyr** | 0.0897 | **231 Ala** | 0.1066 | **361 Gly** | 0.0586 | **415 Gly** | 0.0287 |
| **45 Asp** | 0.0000 | **100 Arg** | 0.0356 | **232 Val** | 0.2493 | **362 Leu** | 0.0058 | **416 Ala** | 0.0261 |
| **46 Gly** | 0.0435 | **101 Ala** | 0.0783 | **237 Val** | 0.0438 | **363 Met** | 0.0720 | **417 Asp** | 0.0358 |
| **47 His** | 0.0499 | **102 Thr** | 0.0208 | **238 Leu** | 0.1061 | **364 Arg** | 0.0381 | **418 Leu** | 0.0353 |
| **48 Ser** | 0.0466 | **103 Leu** | 0.0532 | **239 Asn** | 0.0315 | **365 Ala** | 0.0465 | **419 Ala** | 0.1336 |
| **49 Gly** | 0.0169 | **104 Asp** | 0.0082 | **241 Gln** | 0.0932 | **366 Gly** | 0.0408 | **421 Ser** | 0.0714 |
| **50 Glu** | 0.0477 | **105 Trp** | 0.1153 | **242 Thr** | 0.0314 | **367 Gly** | 0.0276 | **422 Ala** | 0.0557 |
| **51 Val** | 0.0233 | **107 Ala** | 0.0374 | **243 Asn** | 0.0100 | **368 Leu** | 0.0157 | **423 Ala** | 0.1355 |
| **52 Leu** | 0.0683 | **108 Ala** | 0.0494 | **244 Ala** | 0.0347 | **369 Ile** | 0.0473 | **424 Gly** | 0.0240 |
| **53 Gln** | 0.1104 | **109 Gly** | 0.0187 | **251 Ala** | 0.1162 | **370 Leu** | 0.0178 | **425 Leu** | 0.0713 |
| **54 Met** | 0.0230 | **110 His** | 0.1121 | **252 Val** | 1.6149 | **371 Arg** | 0.0147 | **426 Leu** | 0.0450 |
| **55 Ser** | 0.0774 | **111 Leu** | 0.1057 | **254 Arg** | 0.3063 | **372 Phe** | 0.0079 | **427 Ala** | 0.0160 |
| **56 Leu** | 0.0387 | **112 Ser** | 0.0912 | **255 Ala** | 0.6364 | **373 Glu** | 0.0040 | **428 Ile** | 0.0302 |
| **59 Ala** | 0.0527 | **113 Leu** | 0.0537 | **256 Thr** | 0.4893 | **374 Gly** | 0.0194 | **429 Ser** | 0.0093 |
| **60 Thr** | 0.0334 | **117 Arg** | 0.0292 | **257 Ser** | 0.8236 | **375 Arg** | 0.0153 | **435 Ser** | 0.0765 |
| **61 Phe** | 0.3156 | **118 Val** | 0.1336 | **300 Asp** | 0.1235 | **376 Trp** | 0.0174 | **436 Glu** | 0.0146 |
| **62 Leu** | 0.0056 | **119 Gly** | 0.0468 | **301 Leu** | 0.1354 | **377 Gln** | 0.0328 | **437 Cys** | 0.0163 |
| **63 Gly** | 0.1401 | **143 Ala** | 0.0281 | **302 Arg** | 0.1151 | **378 Thr** | 0.0202 | **438 Leu** | 0.0140 |
| **64 Gln** | 0.1165 | **144 Met** | 0.0269 | **303 Thr** | 0.2959 | **379 Leu** | 0.0057 | **439 Val** | 0.0724 |
| **65 Glu** | 0.1009 | **145 Ser** | 0.0401 | **304 Thr** | 0.0873 | **380 Gly** | 0.0095 | **440 Trp** | 0.0348 |
| **67 Thr** | 0.0277 | **146 Ala** | 0.0373 | **305 Leu** | 0.2289 | **381 Glu** | 0.0345 | **441 Leu** | 0.0422 |
| **68 Val** | 0.1100 | **147 Leu** | 0.1007 | **306 Glu** | 0.0784 | **382 Val** | 0.0037 | **442 Arg** | 0.0291 |
| **69 Leu** | 0.0042 | **148 Glu** | 0.0493 | **307 Tyr** | 0.2703 | **387 Ala** | 0.0357 | **445 Leu** | 0.0282 |
| **70 Arg** | 0.0629 | **149 Ser** | 0.0616 | **308 Leu** | 0.2377 | **388 Val** | 0.0236 | **446 Arg** | 0.0265 |
| **71 Gly** | 0.0354 | **150 Ala** | 0.0307 | **309 Gly** | 0.0663 | **390 Ala** | 0.0265 | **447 Leu** | 0.0126 |
| **72 Gln** | 0.0385 | **159 Val** | 0.1090 | **310 Arg** | 0.1040 | **391 Leu** | 0.0326 | **448 Glu** | 0.1006 |
| **73 Thr** | 0.0676 | **160 Ala** | 0.1387 | **311 Glu** | 0.0509 | **392 Leu** | 0.0820 | **449 Val** | 0.0152 |
| **74 Leu** | 0.0675 | **161 Thr** | 0.0380 | **338 Val** | 0.0780 | **393 Ala** | 0.0749 | **450 Ala** | 0.0243 |
| **75 Ala** | 0.0095 | **162 Gln** | 0.1127 | **339 Ala** | 0.0868 | **394 Trp** | 0.1295 | **495 Arg** | 0.0930 |
| **76 Ala** | 0.0115 | **163 Thr** | 0.0746 | **340 Leu** | 0.0739 | **395 Leu** | 0.0209 | **496 Asp** | 0.0295 |
| **77 Leu** | 0.0372 | **164 Val** | 0.0000 | **341 Ala** | 0.0459 | **396 Glu** | 0.0192 | **497 Thr** | 0.0121 |
| **78 Leu** | 0.0180 | **165 Arg** | 0.0283 | **342 Ala** | 0.0226 | **397 Thr** | 0.0201 | **498 Leu** | 0.0768 |
| **80 Glu** | 0.0218 | **166 Glu** | 0.0256 | **343 Ala** | 0.0837 | **398 Gln** | 0.0281 | **499 Thr** | 0.0546 |
| **81 Gln** | 0.0078 | **167 Leu** | 0.0128 | **344 His** | 0.0243 | **402 Leu** | 0.0124 | **500 Gly** | 0.0238 |
| **82 Trp** | 0.0402 | **218 Arg** | 0.1057 | **349 His** | 0.0065 | **403 Val** | 0.0228 | **501 Ala** | 0.0156 |
| **84 Ala** | 0.0319 | **219 His** | 0.0491 | **350 Asp** | 0.0378 | **404 Gln** | 0.0092 | | |
| **85 Leu** | 0.1377 | **220 Leu** | 0.0097 | **351 Thr** | 0.0703 | **405 Thr** | 0.0672 | | |
| **86 Gln** | 0.0000 | **221 Leu** | 0.0530 | **352 Leu** | 0.0460 | **406 Asp** | 0.0229 | | |
| **87 Ala** | 0.0438 | **222 Arg** | 0.0196 | **353 Ser** | 0.0308 | **407 Ala** | 0.0672 | | |

Figure 7: All experimental Chemical Shift Perturbations listed as a table
